# Social and nutritional factors shape larval aggregation, foraging, and body mass in a polyphagous fly

**DOI:** 10.1101/377986

**Authors:** Juliano Morimoto, Binh Nguyen, Shabnam Tarahi Tabrizi, Fleur Ponton, Phillip W. Taylor

## Abstract

The majority of insect species have a clearly defined larval stage during development. Larval nutrition is crucial for individuals’ growth and development, and larval foraging success often depends on both resource availability and competition for those resources. To date, however, little is known about how these factors interact to shape larval development and behaviour. Here we manipulated the density of larvae of the polyphagous fruit fly pest *Bactrocera tryoni* (‘Queensland fruit fly’), and the diet concentration of patches in a foraging arena to address this gap. Using advanced statistical methods of machine learning and linear regression models, we showed that high larval density results in increased larval aggregation across all diets except in extreme diet dilutions. Larval aggregation was positively associated with larval body mass across all diet concentrations except in extreme diet dilutions where this relationship was reversed. Larvae in low-density arenas also tended to aggregate while those in high-density arenas tended to disperse, an effect that was observed for all diet concentrations. Furthermore, larvae in high-density arenas displayed significant avoidance of low concentration diets – a behaviour that was not observed amongst larvae in low-density arenas. Thus, aggregation can help, rather than hinder, larval growth in high-density environments, and larvae may be better able to explore available nutrition when at high-density than when at low density.

## Introduction

In holometabolous insects, larval foraging behaviour largely determines individual fitness (Chapman, 1998). Poor developmental conditions marked by low resource availability – such as when food is scarce and there is high larval competition – often affects both larval developmental time and body size in adulthood [e.g. ^1-10^]. Adult body size tends to correlate positively with female fecundity as well as male mating performance and reproductive success ^5,11^; accordingly, larval foraging behaviour is under productivity selection in females and sexual selection in males ^11-15^, with profound effects on behavioural and evolutionary processes such as cognitive task performance, survival, reproduction, and ultimately sexual selection and sexual conflict ^6,16-18^.

The quantity of resources in a food patch and the number of competing foragers are important determinants of larval responses to developmental conditions ^19^. To maximize resource acquisition for investment in fitness traits of adulthood ^20,21^, larvae are expected to avoid competition with conspecifics, and to prefer patches of highest resource availability. The rationale for this is simple; if the resources are poor or the number of individuals sharing a finite resource is high, the benefits of foraging on that patch may be outweighed by the potential benefits of leaving that patch to seek resources elsewhere. Thus, the ideal situation may be that in which larvae forage in resource-rich food patches without competition. Research across insect taxa has shown that insect larvae have well-defined optimum diets that sustain development and growth, and produce high quality adults ^22-27^, that an excess of nutrients can be detrimental and even compensated for when larvae have a choice to select their food [e.g. ^28-31^]. For social interactions, however, the rule is far less intuitive. Larval aggregations are common in many insects ^32,33^. Although such social interactions may increase foraging competition, larval aggregations can confer physiological and behavioural benefits that sustain larval growth and development ^34-45^. As a result, larvae may maximize development in a high-quality diet with some degree of social interactions and aggregation, provided that competition is not so high that the benefits of aggregation are negated. For instance, *Drosophila* larvae can benefit from occupying patches that are shared with conspecifics, although the increase in competition can in some cases offset the benefits of social behaviour ^45^ [see also ^46-48^]. This hypothesis is derived from the premise that social and nutritional factors interact to shape larval behaviour and growth during development. To date, however, there have been very few direct empirical tests of this hypothesis.

An early attempt to demonstrate interactions between nutritional and social factors as determinants of larval development showed that, in the gregarious caterpillar *Hemileuca lucina,* social environment interacts with the quality of the food source to determine larval growth at mild temperatures ^37^. This investigation only contrasted caterpillars in solitary and groups of a fixed size (10 individuals), and only investigated development on two related-food sources, young vs. mature leaves of *Spiraea latifolia.* Although providing a useful demonstration of concept, this dichotomous approach – i.e. solitary *vs* groups, young *vs* mature leaves – has limited scope for understanding the interaction between social and nutritional factors driving the ecology of larval development. Other studies have shown the importance of larval aggregation in feeding and growth rates, insect-plant interactions, larval defence against predators, and larval thermoregulation [e.g. ^34-44^]. However, there has been no detailed investigation of how the social and nutritional environments of larvae interact to shape development and performance. Key questions remain unanswered though, as to ‘how does the number of foraging larvae with access to a common resource pool, which increase the potential for social interactions, influence larval aggregation?’; ‘When resource availability decreases, do larvae aggregate to the same extent as to when resources are abundant?’; and ‘What are the implications of density- and diet-dependent larval aggregation to larval growth and foraging behaviour?’

In the present study, we addressed these key questions of the interaction between nutritional and social factors driving larval foraging decisions and performance in the tephritid fruit fly *Bactrocera tryoni* (aka ‘Queensland fruit fly’ or ‘Qfly’). Some tephrtids are highly polyphagous and are amongst the most damaging insect pests of horticulture globally ^49-51^. *Bactrocera tryoni* is able to infest more than 150 different fruits ^49,52^; the wide diversity of fruit that are exploited by *B. tryoni,* and variability of nutrients available in infested fruit, make this species well suited for investigation of larval nutritional ecology. Here we first designed circular foraging arenas containing patches of varying macronutrient concentration, where different densities of larvae were allowed to forage. Larvae foraged freely in choice and no-choice arenas, which allowed us to investigate the diet- and density-dependent effects of larval developmental environment on foraging behaviour and larvae body mass. Using statistical methods of machine learning and linear regression, we tested whether tendency to aggregate and size of aggregations depended on the larval density and diet, by allowing groups of several larval densities to forage in arenas of varying diet concentration within which each arena contained multiple patches of the same diet. We then tested how larval density and aggregation affected larval body mass across different diets. Finally, we investigated how larval density influenced larval foraging decisions when facing choices amongst patches with varying resource availability.

### Predictions

1. Previous studies in other species have shown that larvae prefer to occupy patches that are shared with conspecifics [e.g., ^45^]. Thus, we predicted that an increase in larval density should increase aggregation formation as well as aggregation size amongst diet patches. However, this effect could be diet-dependent, whereby macronutrient-poor diets could support smaller aggregations whereas macronutrient-rich diets would support larger aggregations. As a result, we predicted that aggregates should be smaller in macronutrient-poor diets than in macronutrient-rich diets;
2. In other insects, larval aggregation can facilitate feeding [e.g., ^40^]. We therefore predicted that treatments with high larval aggregations should have larvae with higher body mass. However, macronutrient-poor diet is known to reduce larval body mass (see ‘Introduction’). As a result, we predicted that larval body mass should be lower in macronutrient-poor diets compared with macronutrient-rich diets;

## Materials and Methods

### Fly stock and egg collection

We collected eggs from a laboratory-adapted stock of *B. tryoni* (>17 generations-old). The colony has been maintained in non-overlapping generations in a controlled environment room (humidity 65 ± 5%, temperature 25 ± 0.5°C) with light cycle of 12h light: 0.5h dusk:11h dark: 0.5h dawn). Adults were provided a free-choice diet of hydrolysed yeast (MP Biomedicals, Cat. n° 02103304) and commercial refined sucrose (CSR® White Sugar), while larvae were maintained using the Chang-2006 gel-based diet formulation of Moadeli, et al. ^53^ for the last 7 generations (previously maintained on a carrot-based diet). We collected the eggs in a 300mL semi-transparent white plastic bottle that had numerous perforations of <1mm diameter through which females could insert their ovipositor and deposit eggs. The bottle contained 20mL of water, to maintain high humidity. Eggs were collected for 2h, and were then transferred to larval diet with a soft brush, where eggs were allowed to hatch and larvae to develop until they reached 2^nd^ instars.

### Experimental diets and foraging arena

We used 5 experimental diets that varied in macronutrient (i.e., yeast for protein and sugar for carbohydrate) concentration: our control and reference 100% Chang-2006 gel-based diet, which has proven effective for the larvae of this species ^53^, followed by diets with 80%, 60%, 40%, and 20% macronutrient concentration relative to the control diet (see Supplementary Tables for recipes). 20mL of diet was poured into 90mm diameter Petri dishes and allowed to set. We also prepared an agar solution that contained the same components as the gel diets except that no yeast or sugar was included. 20mL of the agar solution was used to cover 90mm diameter Petri dishes that then served as “foraging arenas”. After setting, five equally spaced holes were made in the agar base of each foraging arena by perforating it with a 25mm diameter plastic tube. The same tube was used to cut discs from the experimental diets. The discs of experimental diets were then deposited – in order or randomly – in the holes that had been cut in the agar base of the foraging arenas (see Fig S1). Because the agar solution did not contain macronutrients, we considered the remaining areas of agar base as ‘no choice’ foraging option. Thus, larvae had a total of 6 options (i.e., 5 experimental diets + agar base). The pH of all experimental diets and the agar base was adjusted to 3.8-4 with citric acid. For the experiment, hydrolyzed yeast and sucrose were obtained from MP Biomedicals (Cat. n° 02103304 and 02902978, respectively), Brewer’s yeast was obtained from Lallemand (Cat n° LBI2250), Nipagin was obtained from Southern Biological (Cat n° MC11.2), and all other chemicals composing the diet (e.g., citric acid [see ^53^]) were obtained from Sigma Aldrich®.

### Experimental procedures and statistical analyses

For all experiments, we placed 2^nd^ instar larvae at the centre of the foraging arena (see Fig S1) that was then covered with the lid to minimize the loss of moisture. To minimize potential for effects of visual cues on larval diet choices, the foraging arenas were placed in a dark room. Foraging arenas were set up at 4 larval densities: 10, 25, 50, and 100 larvae. All larvae were released in the arena simultaneously. We did not observe cannibalism or escapes (larval counts were the same at the beginning and at the end of the experiments). All statistical analyses were performed using R version 3.4.0 and plots were performed using the package ‘ggplot2’ ^54,55^.

### Experiment 1: Larval aggregation

To test effects of density and diet on larval aggregation and growth, for all diets and across all larval densities, we set up foraging arenas that contained 5 food patches of the same diet concentration (e.g., all patches with 100% diets) (see Fig S1). We then numbered the patches, and assessed the number of larvae in each of the diet patches as well as on the agar base at 1h, 2h, 4h, 6h, 8h, and 24h after larvae were placed in the arena. We observed that larvae could move across the diameter of the foraging arena in less than 1min, meaning that the time points used in the experiment were ample to allow larvae to explore the entire foraging arena. Four replicates were set up per larval density per diet *(N* = 80 foraging arenas). After 24h, 3 larvae per diet per larval density per replicate were selected from each foraging arena and weighed on a ME5 Sartorius® scale (0.001g precision) to obtain an estimate of average larval body mass. We tested the effects of larval density, diets, and their interaction, using two-way ANOVA model that included replicate as a covariate. To measure larval aggregation, we calculated an ‘aggregation index’ (*AI*) which was the sum of the absolute residuals of our observed data against the machine learning random predictions of a density-dependent random distribution; the procedure to obtain *AI* was as following:

1. We simulated the choices of larvae in foraging arenas with density 10, 25, 50, 100, and 200 larvae choosing amongst 6 patches, where the larvae were equally likely to display choice for any of the options (i.e., the choices for each patch were displayed with equal probability 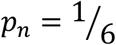, where *p_n_* is the probability of a larvae choosing a given patch). We extrapolated our simulation for larval densities of 10, 25, 50, 100, and 200 larvae in order to build a robust function of density-dependent aggregation (see Fig S2).
2. We then obtained the residual distribution of our empirical data and the simulated density-dependent model against the exact random distribution, calculated simply by dividing the larval density by the number of patch options (i.e. 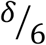, where *δ* = larval density); We then fitted a random forest machine learning regression using the package ‘randomForest’ ^56^ to obtain a model that predicted the behaviour of the residuals. The random forest regression was cross-validated using the package ‘rfUtilities’ ^57^ (Fitted Mean Square Error of the model: 0.009; Median Cross-validation RMSE: 0.036); To build the model, 80% of the simulated data was used in the training phase while 20% was used in the test phase. The model performed accurately during the test phase (Mean Square Error in the Test dataset: 0.038);
3. Next, we used the machine learning model to predict the expected distribution of residuals in our dataset using the ‘predict’ function, and calculated the aggregation index *(AI)* as the difference between the observed sum of residuals and the predicted sum of the residuals obtained with the machine learning regression algorithm.

The machine learning model provides more accurate predictions of the expected distribution of the residuals than conventional linear model. For instance, the MSE (mean square error) of the machine learning model in the test data set was 0.00404 whereas the MSE estimated using conventional linear model was 0.0107, suggesting that the machine learning model was ~2.7 times more accurate in its prediction. We therefore opted to use the machine learning approach to account for non-linear behaviour of the residuals as the density of larvae in the foraging arenas increases (see Fig S2). When we modelled *AI* using general linear model followed by a two-way ANOVA to determine the effect of time, larval density, diet, and their two-way interactions, we transformed *AI* (i. e. *AI*^2.25^) in order to stabilize the variance across larval densities (Levene’s test: F_3,476_ = 0.560, p = 0.641) and diets (Levene’s test: F_4,475_ = 0.548, p = 0.700). To test for the effects of aggregation on larval body mass, we used an ANOVA with the average aggregation index over time, larval density, and diet, as well as the two-way interactions between these factors. For statistical inference, we transformed larval body mass (i.e., *Larval mass* ^0.3^) for homogeneity of variances across larval densities (Levene’s test: F_3,76_ = 0.591, p = 0.622). To calculate the average size of the largest aggregation, we sampled the aggregation with the highest larval count, and calculated the proportion of individuals of the group that were found in that aggregation (ρ) as *ρ* = *α*/*δ*, where α = the number of larvae in the largest patch and δ = the larval density of the group.

To test for the effects of time, larval density, diet, and their two-way interactions we used a generalized linear model (GLM) with *Binomial* distribution – as we were dealing with proportion data – and *quasi* extension, to account for overdispersion of the data. Plots are of the raw data.

### Experiment 2: Larval foraging

For larval foraging assays, the foraging arena contained one patch of each experimental diet, and we assessed the number of larvae selecting each diet across all larval densities (see above) at 1h, 2h, 4h, 6h, and 8h after larvae were placed in the arena. Foraging arenas contained food patches (i.e. 100%, 80%, 60%, 40% and 20% macronutrient concentration) in different orders within the arena (see Fig S1); we controlled for the order of the patches in all models, which had no effect in the results (see ESM). We fitted a multinomial logistic regression model using the ‘multinom’ function of the “nnet” package ^58^. To test for foraging propensity, we controlled for the order of the food patches while investigating the main effects of time, larval density, and their interaction. Agar base (no choice) was the reference level. To test for dietary choices, we used the same multinomial logistic regression, but this time only considering those larvae that chose to forage. By using the standard diet (100% macronutrient concentration) as our reference level, we could then infer the relative dietary preferences of larvae that foraged. Statistical inferences for multinomial logistic regressions were made based on 95% and 99% confidence intervals for each larval density separately.

## Results

### Experiment 1: High larval density increases larval body mass

We first tested the influence of larval density on growth. Our results showed highly significant positive effects of diet concentration and larval density on body mass (Table S1), although there was no significant interaction between these factors. Body mass increased steadily with larval density in the foraging arena and consistently across all diets (Fig 1). However, diet concentration also affected larval body mass, as larvae from foraging arenas with diluted diets (i.e. 40% and 20% macronutrient concentration) had lower body mass than larvae from arenas containing more concentrated diets (Fig 1).

**Figure 1.**
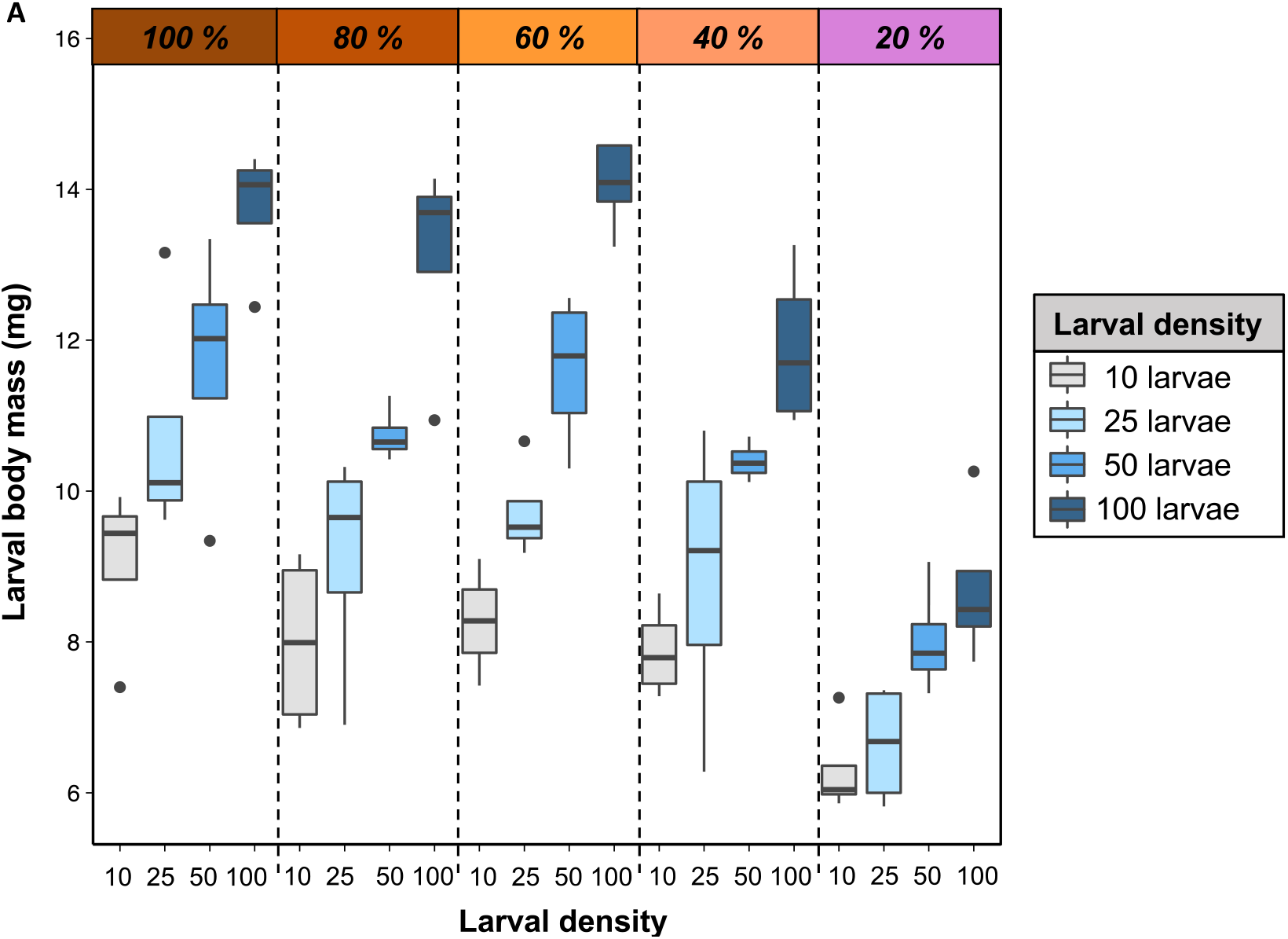
Schemes of Myo1b-Actin gliding assays. Larval density increases larval body mass across diet dilutions. Body mass (mg) of larvae from different larval densities and from across diets, at the end of our experiment (24h, see Methods for details).

### Experiment 1: Larval density affects larval aggregation in a diet-dependent manner

We investigated whether larval density modulated larval aggregation, and whether this relationship was affected by diet concentration. We found significant interaction between effects of diet concentration and larval density on the aggregation index (Table S2), whereby larvae in high-density arenas aggregated more in high macronutrient concentration diets (>40%) and less in low macronutrient concentration diets (20%, Fig 2a-b).

**Figure 2.**
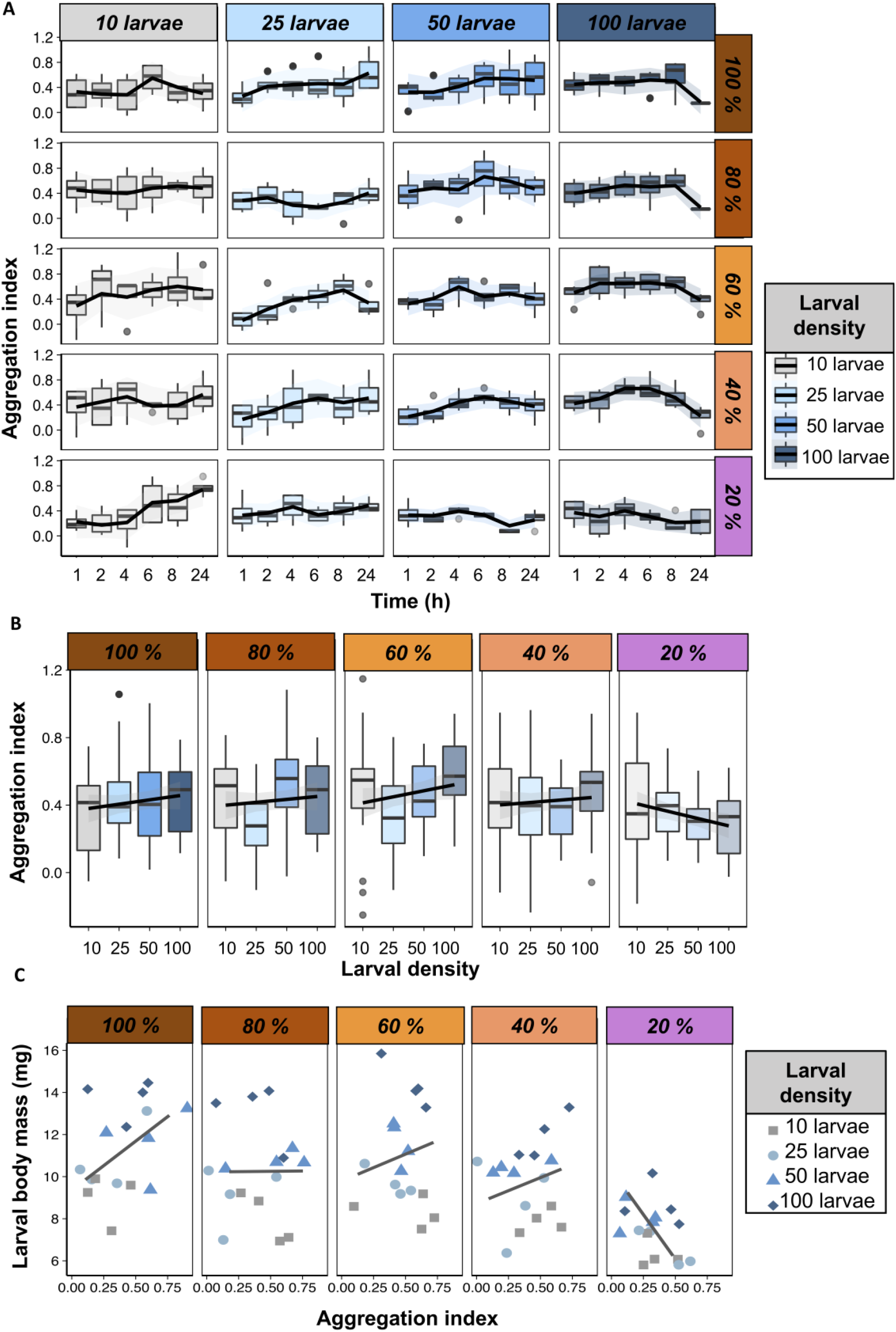
Body mass and the relationship between body mass and aggregation. (a) Larval aggregation index (*y-*axis) over time (*x-*axis) across larval densities (horizontally) and across diets (vertically). Lines were drawn using the ‘loess’ method in the package ‘ggplot2’ in R, and indicate the trend in the data. (b) Average larval aggregation index (y-axis) on larval density (x-axis) over all time points in our experiment. Lines were drawn using the ‘lm’ method in the package ‘ggplot2’ in R, and indicate the trend in the data. (c) The relationship between larval body mass and the average aggregation index. Colours and shapes indicate the larval density. Lines were drawn using the ‘loess’ method in the package ‘ggplot2’ in R, and indicate the trend in the data.

There was a significant interaction between time and larval density, whereby larvae in low-density arenas (10 larvae) increased aggregation as time foraging passed, while the opposite pattern was observed for high-density arenas (100 larvae) (Table S2, Fig 2a-b). This was particularly evident for low-density arenas with low macronutrient concentration diets (see Fig 2a). This is important because if the larvae were simply coalescing in the same location (i.e., not seeking to aggregate but converging to the same location with high quality food substrate), we would expect larvae in low-density arenas to show the same pattern for high- and low diet concentration. Instead, the results show the opposite is true, whereby larvae in low-density arenas tended to aggregate more over time with low diet concentration than with high diet concentration (Fig 2a). This provides evidence that larvae seek to aggregate, especially when foraging in low-density arena and with low-resource food substrates. Arenas with density of 25 and 50 larvae showed the same trend as arenas with 10 and 100 larvae, respectively, although with lower magnitude (Fig 2a-b).

### Experiment 1: The relationship between larval aggregation and larval body mass is diet-dependent

Next, we tested the relationship between larval aggregation and body mass. We found that aggregation had an overall highly significant positive effect on larval body mass when diet concentration was 40% or greater but that a negative trend was instead observed when diet concentration was 20% (Fig 2c, Table S3). There was a significant effect of diet concentration and larval density, but there were no significant interactions between larval density and diet concentration, larval density and aggregation index, nor between diet concentration and aggregation index (Table S3). These results provide evidence for a positive relationship between larval aggregation and larval body mass, and revealed that in some cases nutrient concentration in the diet can be a strong modulator of this relationship.

### Experiment 1: Larval density and diet influence the size of larval aggregations

Previous studies have shown that larval aggregation can help larvae to feed more efficiently, potentially leading to an increase in larval body mass (see for instance ^40,59^). If this is true, an aggregation could become a ‘hotspot’ for other larvae, and we would expect that arenas with high larval densities would have few large aggregations. This could explain the relationships between larval aggregation and body mass and also the relationship between larval density and larval aggregation. Alternatively, high larval density could make larvae more inclined to disperse in order to minimize competition and, as a result, form smaller aggregations at more locations, hence exploiting a greater number of food patches. Our results showed a significant interaction between the effects of larval density and time, and larval density and diet concentration on the proportion of individuals in the largest aggregation (see Table S4, Fig 3). These results demonstrate that i) arenas containing diluted diets (i.e., 20% and 40%) had relatively more larvae in the largest aggregations than did arenas containing more concentrated diets, ii) low larval density arenas (i.e., 10 larvae) had aggregations that contained relatively more larvae compared with higher density arenas (i.e., 25, 50, 100), iii) high density arenas (i.e., 100 larvae) were more evenly distributed compared with low density arenas, whereas the opposite effect was found for low density arenas (i.e., 10 larvae), and iv) the proportion of larvae in the most numerous aggregation decreased in diluted diets in high density arenas, an effect that was not observed for low-density arenas (see Fig 3). These findings support the hypothesis that high larval density promotes larval movement, whereby larvae formed smaller aggregations that exploit patches more evenly.

**Figure 3.**
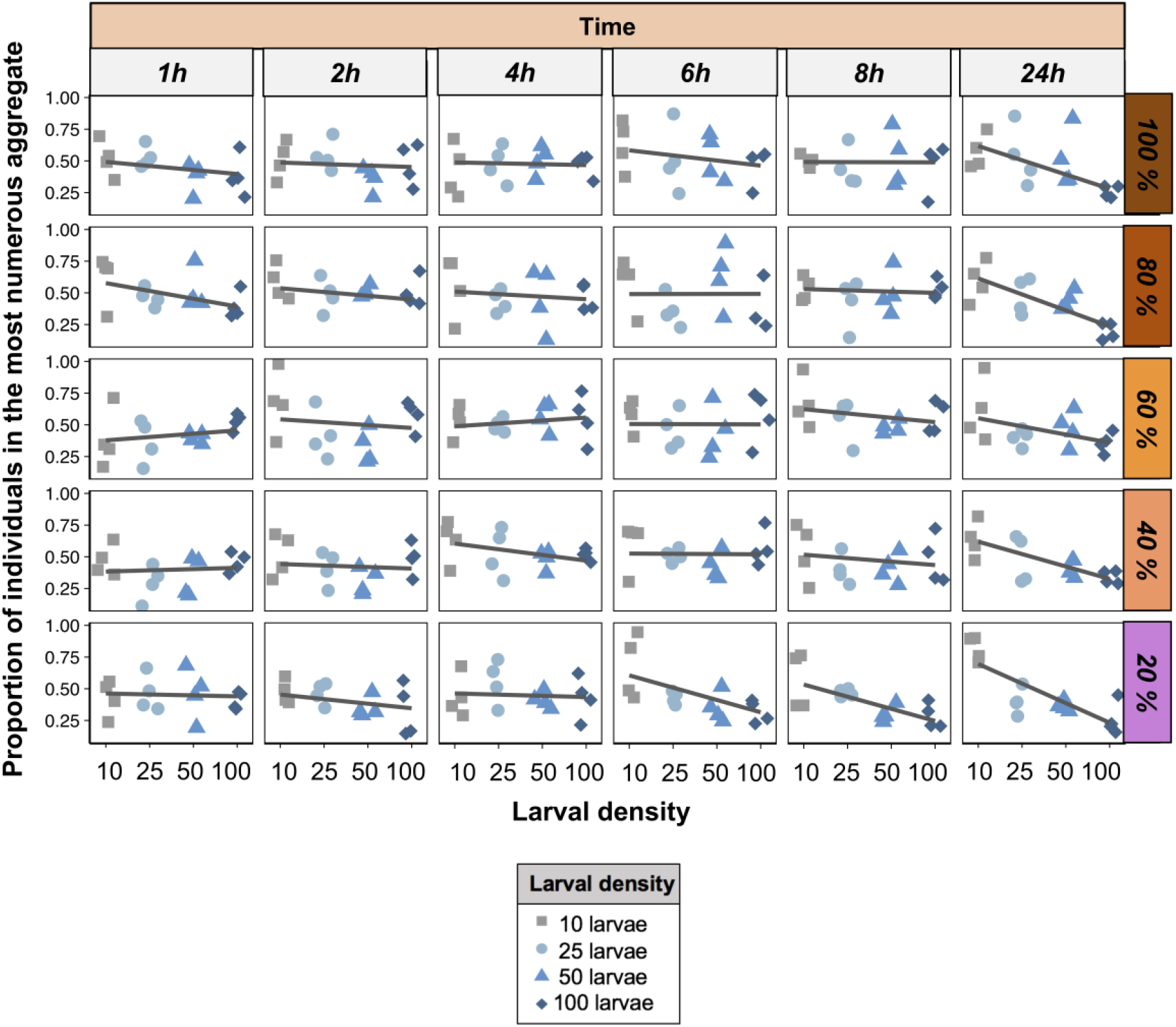
Body mass and the relationship between body mass and aggregation. Proportion of larvae in aggregates. The proportion of individuals in the most numerous aggregate over time (horizontally) across diets (vertically). Shapes and colours indicate larval density. Lines were drawn using the ‘lm’ method in the package ‘ggplot2’ in R, and indicate the trend in the data.

### Experiment 2: Larval density shapes larval foraging behaviour

Next, we measured how larval density influenced larvae foraging propensity, as well as larvae foraging decisions when larvae have a choice amongst patches with varying diet concentrations. By using a multinomial logistic regression model that used ‘no choice’ (i.e. agar base) as our reference level, we could assess larval foraging propensity over time. Our results showed that larvae were more likely to forage in any given patch than to not forage at all, and the propensity of foraging was particularly high for patches of high nutrient concentration independent of larval density (Fig 4a, Table S5, Fig S3). Interestingly, the range of diets in which larvae foraged was greater for arenas containing 50 and 100 larvae and included the patch with 40% diet in addition to the 100%, 80% 60% patches that were more dominant for arenas of lower larval density (Fig 4a). These findings show that larvae are generally more prone to forage in high-quality patches, and that larval foraging propensity is density-independent.

**Figure 4.**
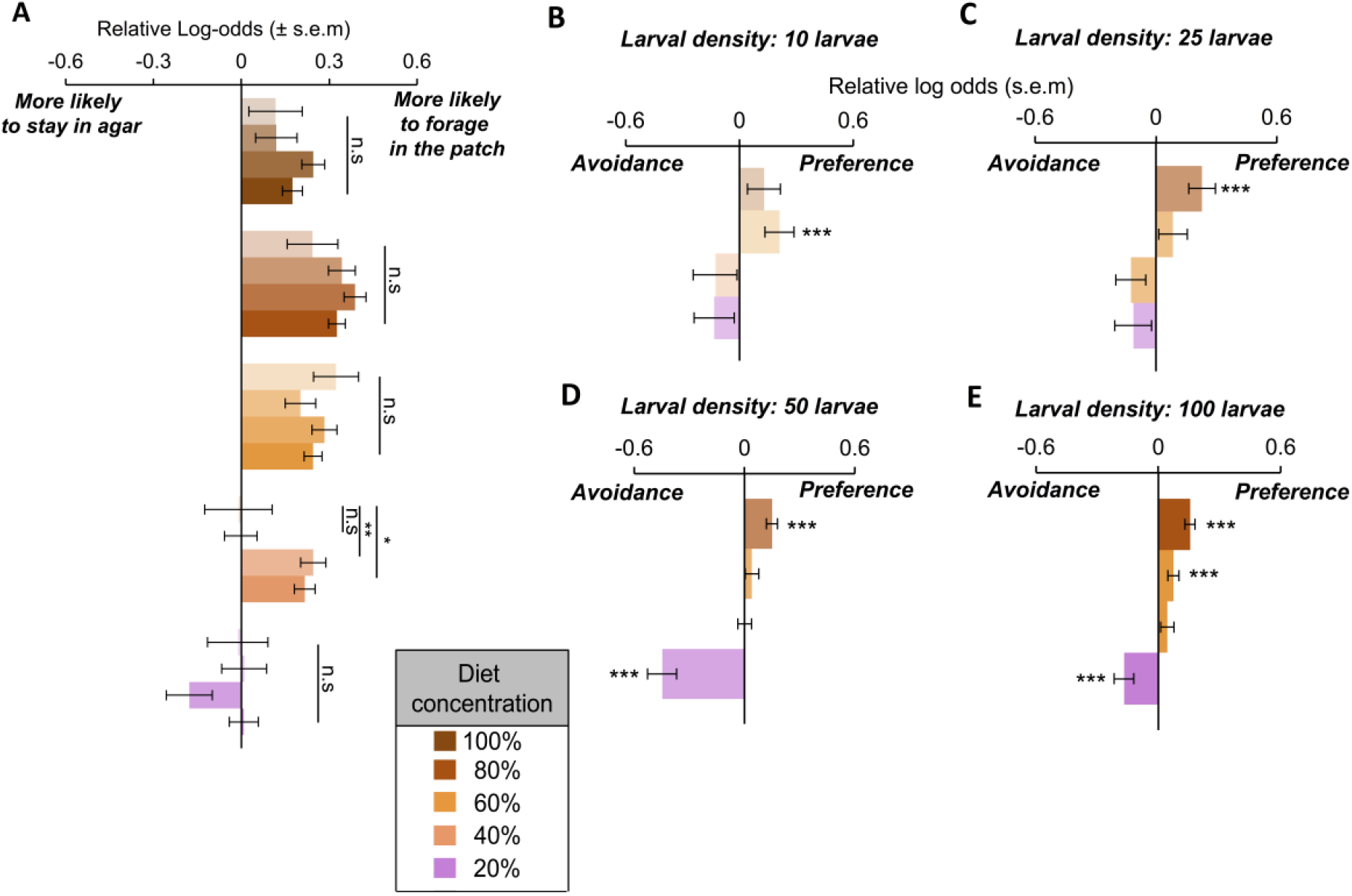
Larval foraging propensity. (a) Relative log-odds of larvae making a choice to forage in a given food patch relative to staying in agar (no choice). Shades represent different larval densities: 10, 25, 50, and 100 larvae. p-values obtained using Students’ t-distribution. Note that relative log-odds are calculated using the control 100% diet as reference. Log-odds > 0: more likely to choose a given patch relative to staying in agar, Log-odds < 0, less likely to choose a given patch relative to staying in agar. s.e.m = standard error of the mean. (b-e) Relative log-odds of larvae patch preferences. Patch with standard diet (100% macronutrient concentration) was used as the reference level. *** non-overlapping 99% confidence intervals. s.e.m = standard error of the mean.

We then tested whether larval density affected larval diet choices, using again a multinomial logistic regression although this time we used the standard diet (i.e. 100%) as our reference diet and excluded non-foraging larvae, while modelling the behaviour of larvae that were actively foraging in one of the food patches in the previous experiment. In arenas with low larval density (10 larvae), larvae displayed a significant preference for diets with 60% macronutrient concentration relative to the standard (100%) diet (Fig 4b, Table S6). However, as larval density increased (25 and 50 larvae), there was a shift in preference toward the patch containing 80% macronutrient concentration (Fig 4b), and finally, when larval density was the highest (100 larvae), larvae displayed statistically significant preferences for both 60% and 80% macronutrient patches compared to the standard diet (Fig 4b-e). More importantly, though, is that only larvae in arenas with high density (50 and 100 larvae) displayed significant avoidance of low concentration patches of 20% macronutrient concentration (Fig 4d-e).

## Discussion

In this study, we demonstrate how key ecological factors interact to determine larval foraging behaviour and growth in *B. tryoni*. Our findings showed that larval aggregation increased with larval density in a diet-dependent manner, and promoted larval body mass across all diets. Importantly, larval density modulated the size of larval aggregations, and influenced larval foraging behaviour when larvae experienced patches with varying concentrations, highlighting a role of social interactions and population density for larval behaviour. Our findings provide insight into larval foraging decisions of fruit flies, and more generally, provide insight into broad ecological patterns arising from nutrition and intraspecific competition within groups and populations. Fruit fly larvae are commonly found in aggregations within a fruit ^9,10,60^. Furthermore, fruits can be heterogeneous foraging environments for larvae [e.g., ^61^], and the nutritional composition of fruit can change as larvae develop [see ^62-64^]. Therefore, the density of larvae and local diet quality might determine larval movement within a fruit in search of more nutritious and less competitive foraging sites. It is important to note that it is unlikely that our findings apply to movement of larvae between fruits. Crawling out of fruits is dangerous owing to risks of predation [^65^, reviewed by ^66^] and desiccation. In nature, *B. tryoni* females modulate their oviposition behaviour to minimize intra-specific competition amongst larvae ^67^, and it is reasonable to expect the larvae to very rarely move between fruits.

High population density can force animals to change their behaviour and expand their niche due to inter- and intra-specific competition, and this is a well-established ecological principle observed in both the laboratory and in nature ^68,69^. Even though larvae are prone to aggregate, an increase in larval density could increase larval competition within large aggregations, which could in turn drive larvae to disperse and form smaller aggregations across different locations. The smaller aggregation size observed in high-density arenas support this idea, meaning that larval aggregations formed in high-density arenas were proportionally smaller than those formed at lower densities. Moreover, larval aggregations were proportionally smaller as the density increased and the larvae spent more time foraging, suggesting that social interactions within larger aggregations are likely to induce more frequent movement by the larvae. As the larvae move more often, they are more likely to find new (and unexplored) food patches, and are therefore more likely to explore patches more evenly. The influence of larval density on larval aggregation and growth could therefore be a plastic response to intraspecific competition because it could lead to better larval foraging decisions and a broader niche exploration ^45,70^. The findings that high larval density also influence larval foraging behaviour in ways that decrease larval foraging propensity on resource-poor diet patches provide further support for the idea that high larval density promotes exploration of the foraging environment and effective exploitation of nutritional resources. Individuals of many species use social cues when making decisions ^71^, and recent models have predicted that social interactions could improve individual foraging success, especially when food is scarce and distributed heterogeneously ^72^. It is also possible that larval aggregation alters the nutritional composition and the microbial communities of the diets. For instance, larvae of some insect species can be cannibalistic ^73,74^, and because larvae are a rich source of nutrients, cannibalism could affect the nutrient status of a food patch. Moreover, in *D. melanogaster,* larval foraging behaviour is determined by the bacterial communities in the diet ^75^, and in *B. tryoni,* gut-microbial fungi in the diet have been found to promote larval development under nutrient-limiting conditions ^76^. If larval density affected the relative abundance of these fungi in the diet, this could in turn have influenced larval foraging behaviour and larval body mass. Future studies that investigate the impact of larval density on the occurrence of cannibalism, and that compare changes in larval and diet microbial profiles in high- and low-density social environments will provide insights into the mechanisms underpinning the effects of larval environments on foraging behaviour and growth.

A negative relationship between population density and individual fitness is often assumed in ecology [reviewed by ^77^]. In invertebrates, including tephritid fruit flies, high-densities at the larval stage can decrease nutrient availability, and reduce adult body mass, reproductive success, and survival [e.g. ^1,3-9,60^], which can lead to a density-dependent effects on fitness that extends through generations ^6^. However, high densities can also mitigate the negative effects of environmental stresses, and act as a buffering factor for individual fitness and survival [reviewed by ^77^]. Therefore, high-density environments can sometimes confer fitness benefits. Our findings support this view, as they reveal that the density of larvae can trigger behavioural responses early in life that can benefit larval growth. This positive effect is likely due to an increase in exploratory behaviour when at high-densities, which can increase niche exploration and nutrient acquisition. It is important to mention that competition amongst conspecifics should determine threshold in which sociality provides benefits to the larvae, after which further increase in density should incur costs that offset the benefits to individuals’ fitness ^45^. This threshold is currently unknown, but we predict that further increase in the density of larvae in our experiments (e.g., 400 larvae) should result in measurable costs such as decrease in body mass of the larvae. Determining the threshold is out of the scope of this study, but remains an important topic for future investigations. Nonetheless, our findings are applicable to biological scenarios where intraspecific competition increases and resources are heterogeneous, and thus represent a logical consequence of the interaction between the nutritional and social environments.

It is important to mention that as density increases, larvae may be displaced from the patch due to the competition with conspecifics for space. This is a natural consequence of high larval density (i.e., defined as more larvae per unit of space), and understanding how the competition for space underlies larval behaviour is out of the scope of this study. Also, patch quality could have decreased over time, especially in treatments with high larval densities, and influenced some of the results found in our study. This is unlikely, however, because the number of individuals in each patch sharply increased and stabilised in a plateau, with no evidence of larvae evasion from the chosen patches throughout the 24h in which the experiment was conducted (see e.g., Fig S3). Thus, our results demonstrate how the interactions between larval density and larval nutritional environment shape larval foraging behaviour.

## Conclusion

The present study provides a new perspective on density-dependent effects on larval development. Fruit fly larvae respond to a range of social and nutritional factors, with important implications for larval foraging and growth. Together, our findings help us understand the ecological factors underpinning larval development in insects, and serve as an important stepping-stone for future research aimed at better understanding the behavioural and nutritional aspects of development in group-living insects.

## Acknowledgements

Project *Larval diets for high-productivity mass-rearing* (HG13045) is funded by the Hort Frontiers Fruit Fly Fund, part of the Hort Frontiers strategic partnership initiative developed by Hort Innovation, with co-investment from Macquarie University and contributions from the Australian Government. We acknowledge Dr Alistair M Senior, University of Sydney, for helpful comments on the statistical analysis using multinomial logistic regression.

## Authors’ Contribution

J.M, P.W.T, and F.P designed the experiment. J.M, F.P., B.N and S.T.T collected the data. J.M, F.P, B.N and P.W.T analysed the data. All authors wrote the manuscript, approved the submission, and agree to be accountable for all aspects of the work.

## Supplementary Information

**Supplementary Information file** – Supplementary figures and tables with the complete outputs of the statistical models.

**Figure S1 – Design of the foraging arenas.** (a) Schematic representations of the foraging arena used in our larval dietary choice experiments. (b) Schematic representations of the foraging arena used in our larval aggregation experiments. Note that the arenas were designed exactly as in (a), although all patches contained the same diet concentration.

**Figure S2 – Box plots showing the behaviour of the residuals from the density-dependent simulation.** Note that we extrapolated our simulations to include foraging groups with density of 200 larvae (see Methods). Line was drawn using the ‘loess’ method in the package ‘ggplot2’ in R to highlight the trend in the data.

**Figure S3 – Larval foraging behaviour.** The number of larvae in each foraging patch over time across the larval density treatments.

**Table S1 – Complete analysis of larvae body mass. Bold:** p <0.05

**Table S2 – Complete analysis of larvae aggregation index.** Data was transformed (square-rooted) for statistical testing. **Bold:** p <0.05

**Table S3 – Complete analysis of the relationship between larvae body mass and aggregation index. Bold:** p <0.05

**Table S4 – Complete analysis of the proportion of larvae in the most numerous aggregate.** GLM with Binomial distribution and quasi extension to account for overdispersion of the data.

**Table S5 - Complete analysis of larvae willingness to forage.** Agar (no choice) is the reference level. **Bold:** p <0.05

**Table S6 - Complete analysis of larvae dietary choices.** Standard diet (100% macronutrient concentration) as reference level. *** non-overlapping 99% CI.

**Diet recipes** – The formulations for the diets used in the experiments.

## References

1 Amitin, E. G. & Pitnick, S. Influence of developmental environment on male- and female-mediated sperm precedence in *Drosophila melanogaster*. J Evolution Biol 20, 381–391, doi:Doi 10.1111/J.1420-9101.2006.01184.X (2007).

2 Pitnick, S. & Garcia-Gonzalez, F. Harm to females increases with male body size in *Drosophila melanogaster*. P Roy Soc B-Biol Sci 269, 1821–1828, doi:Doi10.1098/Rspb.2002.2090 (2002).

3 Lyimo, E., Takken, W. & Koella, J. Effect of rearing temperature and larval density on larval survival, age at pupation and adult size of Anopheles gambiae. Entomologia experimentalis et applicata 63, 265–271 (1992).

4 Credland, P. F., Dick, K. M. & Wright, A. W. Relationships between larval density, adult size and egg production in the cowpea seed beetle, *Callosobruchus maculatus*. Ecological Entomology 11, 41–50, doi:10.1111/j.1365-2311.1986.tb00278.x (1986).

5 Wigby, S., Perry, J. C., Kim, Y. H. & Sirot, L. K. Developmental environment mediates male seminal protein investment in *Drosophila melanogaster*. Funct Ecol 30, 410–419 (2015).

6 Morimoto, J., Ponton, F., Tychsen, I., Cassar, J. & Wigby, S. Interactions between the developmental and adult social environments mediate group dynamics and offspring traits in *Drosophila melanogaster*. Scientific Reports 7, 3574, doi:10.1038/s41598-017-03505-2 (2017).

7 Moczek, A. P. Horn polyphenism in the beetle *Onthophagus taurus*: larval diet quality and plasticity in parental investment determine adult body size and male horn morphology. Behavioral Ecology 9, 636–641 (1998).

8 Nasci, R. S. & Mitchell, C. J. Larval diet, adult size, and susceptibility of *Aedes aegypti* (Diptera: Culicidae) to infection with Ross River virus. Journal of Medical Entomology 31, 123–126 (1994).

9 Burrack, H. J. et al. Intraspecific larval competition in the olive fruit fly (Diptera: Tephritidae). Environmental entomology 38, 1400–1410 (2009).

10 Averill, A. L. & Prokopy, R. J. Intraspecific competition in the tephritid fruit fly *Rhagoletis pomonella*. Ecology 68, 878–886 (1987).

11 Bonduriansky, R. The evolution of male mate choice in insects: a synthesis of ideas and evidence. Biol Rev 76, 305–339 (2001).

12 Honek, A. Intraspecific variation in body size and fecundity in Insects - a general relationship. Oikos 66, 483–492, doi:Doi 10.2307/3544943 (1993).

13 Roff, D. A. Life history evolution. Vol. 7 (Sinauer Associates Sunderland, 2002).

14 Stearns, S. C. The evolution of life histories. Vol. 249 (Oxford University Press Oxford, 1992).

15 Clutton-Brock, T. Sexual selection in females. Anim Behav 77, 3–11 (2009).

16 Sokolowski, M. B., Kent, C. & Wong, J. *Drosophila* larval foraging behaviour: developmental stages. Anim Behav 32, 645–651 (1984).

17 Sokolowski, M. B., Pereira, H. S. & Hughes, K. Evolution of foraging behavior in *Drosophila* by density-dependent selection. Proceedings of the National Academy of Sciences 94, 7373–7377 (1997).

18 Kohn, N. R. et al. Social environment influences performance in a cognitive task in natural variants of the foraging gene. PloS one 8, e81272 (2013).

19 Bell, W. J. Searching behaviour: the behavioural ecology of finding resources. (Springer Science & Business Media, 2012).

20 Rowe, L. & Houle, D. The lek paradox and the capture of genetic variance by condition dependent traits. P Roy Soc B-Biol Sci 263, 1415–1421, doi:Doi 10.1098/Rspb.1996.0207 (1996).

21 Hill, G. E. Condition-dependent traits as signals of the functionality of vital cellular processes. Ecol Lett 14, 625–634, doi:Doi 10.1111/J.1461-0248.2011.01622.X (2011).

22 Simpson, S. J. & Raubenheimer, D. The nature of nutrition: a unifying framework from animal adaptation to human obesity. (Princeton University Press, 2012).

23 Rodrigues, M. A. et al. *Drosophila melanogaster* larvae make nutritional choices that minimize developmental time. Journal of insect physiology 81, 69–80 (2015).

24 Zucoloto, F. S. Feeding habits of *Ceratitis capitata* (Diptera: Tephritidae): can larvae recognize a nutritionally effective diet? Journal of Insect Physiology 33, 349–353 (1987).

25 Zucoloto, F. Effects of flavour and nutritional value on diet selection by *Ceratitis capitata* larvae (Diptera, Tephritidae). Journal of Insect Physiology 37, 21–25 (1991).

26 Silva-Soares, N. F., Nogueira-Alves, A., Beldade, P. & Mirth, C. K. Adaptation to new nutritional environments: larval performance, foraging decisions, and adult oviposition choices in *Drosophila suzukii*. BMC ecology 17, 21 (2017).

27 de Carvalho, M. J. A. & Mirth, C. K. Food intake and food choice are altered by the developmental transition at critical weight in *Drosophila melanogaster*. Anim Behav 126, 195–208 (2017).

28 Fanson, B. G. & Taylor, P. W. Protein: carbohydrate ratios explain life span patterns found in Queensland fruit fly on diets varying in yeast: sugar ratios. Age 34, 1361–1368 (2012).

29 Um, S. H., D’Alessio, D. & Thomas, G. Nutrient overload, insulin resistance, and ribosomal protein S6 kinase 1, S6K1. Cell Metab 3, 393–402, doi:Doi 10.1016/J.Cmet.2006.05.003 (2006).

30 Musselman, L. P. et al. A high-sugar diet produces obesity and insulin resistance in wild-type Drosophila. Disease models & mechanisms 4, 842–849, doi:10.1242/dmm.007948 (2011).

31 Schwarz, S., Durisko, Z. & Dukas, R. Food selection in larval fruit flies: dynamics and effects on larval development. Naturwissenschaften 101, 61–68 (2014).

32 Taylor, L. Aggregation, variance and the mean. Nature 189, 732–735 (1961).

33 Taylor, L., Woiwod, I. & Perry, J. The density-dependence of spatial behaviour and the rarity of randomness. The Journal of Animal Ecology, 383–406 (1978).

34 Cornell, J. C., Stamp, N. E. & Bowers, M. D. Developmental change in aggregation, defense and escape behavior of buckmoth caterpillars, *Hemileuca lucina* (Saturniidae). Behavioral Ecology and Sociobiology 20, 383–388 (1987).

35 Klok, C. & Chown, S. Assessing the benefits of aggregation: thermal biology and water relations of anomalous Emperor Moth caterpillars. Functional ecology 13, 417–427 (1999).

36 Slone, D. & Gruner, S. V. Thermoregulation in larval aggregations of carrion-feeding blow flies (Diptera: Calliphoridae). Journal of Medical Entomology 44, 516–523 (2007).

37 Stamp, N. E. & Bowers, M. D. Variation in food quality and temperature constrain foraging of gregarious caterpillars. Ecology 71, 1031–1039 (1990).

38 Wise, M. J., Kieffer, D. L. & Abrahamson, W. G. Costs and benefits of gregarious feeding in the meadow spittlebug, *Philaenus spumarius*. Ecological Entomology 31, 548–555 (2006).

39 Hambäck, P. Density-dependent processes in leaf beetles feeding on purple loosestrife: aggregative behaviour affecting individual growth rates. Bulletin of entomological research 100, 605–611 (2010).

40 Denno, R. & Benrey, B. Aggregation facilitates larval growth in the neotropical nymphalid butterfly *Chlosyne janais*. Ecological Entomology 22, 133–141 (1997).

41 Storer, A., Wainhouse, D. & Speight, M. The effect of larval aggregation behaviour on larval growth of the spruce bark beetle *Dendroctonus micans*. Ecological Entomology 22, 109–115 (1997).

42 Brian, R. The consequences of larval aggregation in the butterfly *Chlosyne lacinia*. Ecological Entomology 22, 408–415 (1997).

43 Hochuli, D. F. Insect herbivory and ontogeny: How do growth and development influence feeding behaviour, morphology and host use? Austral Ecology 26, 563–570 (2001).

44 Hunter, A. F. Gregariousness and repellent defences in the survival of phytophagous insects. Oikos 91, 213–224 (2000).

45 Durisko, Z. & Dukas, R. Attraction to and learning from social cues in fruitfly larvae. Proceedings of the Royal Society of London B: Biological Sciences 280, 20131398 (2013).

46 Golden, S. & Dukas, R. The value of patch-choice copying in fruit flies. PloS one 9, e112381 (2014).

47 Venu, I., Durisko, Z., Xu, J. & Dukas, R. Social attraction mediated by fruit flies’ microbiome. Journal of Experimental Biology 217, 1346–1352 (2014).

48 Ward, A. & Webster, M. Sociality: the behaviour of group-living animals. (Springer, 2016).

49 Clarke, A. R., Powell, K. S., Weldon, C. W. & Taylor, P. W. The ecology of *Bactrocera tryoni* (Diptera: Tephritidae): what do we know to assist pest management? Annals of Applied Biology 158, 26–54 (2011).

50 Virgilio, M., Delatte, H., Backeljau, T. & De Meyer, M. Macrogeographic population structuring in the cosmopolitan agricultural pest *Bactrocera cucurbitae* (Diptera: Tephritidae). Molecular Ecology 19, 2713–2724 (2010).

51 Malacrida, A. et al. Globalization and fruitfly invasion and expansion: the medfly paradigm. Genetica 131, 1 (2007).

52 Schutze, M. K., Virgilio, M., Norrbom, A. & Clarke, A. R. Tephritid integrative taxonomy: Where we are now, with a focus on the resolution of three tropical fruit fly species complexes. Annual review of entomology 62, 147–164 (2017).

53 Moadeli, T., Taylor, P. W. & Ponton, F. High productivity gel diets for rearing of Queensland fruit fly, *Bactrocera tryoni*. Journal of Pest Science 2, 507–520 (2017).

54 R Development Core Team. R: A language and environment for statistical computing. R. Foundation for Statistical Computing, Vienna, Austria. http://www.r-project.org/ (2017).

55 Wickham, H. ggplot2: elegant graphics for data analysis. (2009).

56 Liaw, A. & Wiener, M. Classification and regression by randomForest. R news 2, 18–22 (2002).

57 Murphy, M. A., Evans, J. S. & Storfer, A. Quantifying *Bufo boreas* connectivity in Yellowstone National Park with landscape genetics. Ecology 91, 252–261 (2010).

58 Ripley, B. D. Modern applied statistics with S. (Springer, 2002).

59 Prokopy, R. J. & Roitberg, B. D. Joining and avoidance behavior in nonsocial insects. Annual review of entomology 46, 631–665 (2001).

60 Ekesi, S., Billah, M. K., Nderitu, P. W., Lux, S. A. & Rwomushana, I. Evidence for competitive displacement of *Ceratitis cosyra* by the invasive fruit fly *Bactrocera invadens* (Diptera: Tephritidae) on mango and mechanisms contributing to the displacement. Journal of Economic Entomology 102, 981–991 (2009).

61 Peiris, K., Dull, G., Leffler, R. & Kays, S. Near-infrared spectrometric method for nondestructive determination of soluble solids content of peaches. Journal of the American Society for Horticultural Science 123, 898–905 (1998).

62 Matavelli, C., Carvalho, M. J. A., Martins, N. E. & Mirth, C. K. Differences in larval nutritional requirements and female oviposition preference reflect the order of fruit colonization of *Zaprionus indianus* and *Drosophila simulans*. Journal of insect physiology 82, 66–74 (2015).

63 Drew, R. Amino acid increases in fruit infested by fruit flies of the family Tephritidae. Zoological Journal of the Linnean Society 93, 107–112 (1988).

64 MacCollom, G., Lauzon, C., Sjogren, R., Meyer, W. & Olday, F. Association and attraction of blueberry maggot fly Curran (Diptera: Tephritidae) to *Pantoea* (Enterobacter) *agglomerans*. Environmental Entomology 38, 116–120 (2009).

65 Aluja, M., Sivinski, J., Rull, J. & Hodgson, P. J. Behavior and predation of fruit fly larvae (*Anastrepha spp*.)(Diptera: Tephritidae) after exiting fruit in four types of habitats in tropical Veracruz, Mexico. Environmental Entomology 34, 1507–1516 (2005).

66 Uchôa, M. in Integrated Pest Management and Pest Control-Current and Future Tactics (InTech, 2012).

67 Fitt, G. P. Oviposition behaviour of two tephritid fruit flies, *Dacus tryoni* and *Dacus jarvisi*, as influenced by the presence of larvae in the host fruit. Oecologia 62, 37–46 (1984).

68 Svanbäck, R. & Bolnick, D. I. Intraspecific competition drives increased resource use diversity within a natural population. Proceedings of the Royal Society of London B: Biological Sciences 274, 839–844 (2007).

69 Svanbäck, R. & Bolnick, D. I. Intraspecific competition affects the strength of individual specialization: an optimal diet theory method. Evolutionary Ecology Research 7, 993–1012 (2005).

70 Bolnick, D. I. et al. The ecology of individuals: incidence and implications of individual specialization. The American Naturalist 161, 1–28 (2002).

71 Dall, S. R., Giraldeau, L.-A., Olsson, O., McNamara, J. M. & Stephens, D. W. Information and its use by animals in evolutionary ecology. Trends Ecol Evol 20, 187–193 (2005).

72 Lihoreau, M. et al. Collective foraging in spatially complex nutritional environments. Philosophical Transactions of the Royal Society B: Biological Sciences 372, 20160238 (2017).

73 Vijendravarma, R. K., Narasimha, S. & Kawecki, T. J. Predatory cannibalism in *Drosophila melanogaster* larvae. Nature communications 4, 1789 (2013).

74 Schultner, E., d’Ettorre, P. & Helanterä, H. Social conflict in ant larvae: egg cannibalism occurs mainly in males and larvae prefer alien eggs. Behavioral Ecology 24, 1306–1311 (2013).

75 Wong, A. C.-N. et al. Gut microbiota modifies olfactory-guided microbial preferences and foraging decisions in *Drosophila*. Current Biology 27, 2397–2404. e2394 (2017).

76 Piper, A. M., Farnier, K., Linder, T., Speight, R. & Cunningham, J. P. Two gut-associated yeasts in a Tephritid fruit fly have contrasting effects on adult attraction and larval survival. Journal of Chemical Ecology 43, 891–901 (2017).

77 Bruno, J. F., Stachowicz, J. J. & Bertness, M. D. Inclusion of facilitation into ecological theory. Trends in Ecology & Evolution 18, 119–125 (2003).

